# Dynein contributes to spindle elongation during anaphase in *Candida albicans*

**DOI:** 10.1101/2024.08.15.608126

**Authors:** Irsa Shoukat, John S. Allingham

## Abstract

Budding yeast must position the mitotic spindle at the mother-bud junction during anaphase to ensure proper chromosome segregation. Pushing and pulling forces on astral microtubules from dynein motors in the cell cortex are crucial for spindle positioning. In some higher eukaryotes and filamentous fungi, dynein also accelerates spindle elongation during anaphase, but in yeast model systems spindle elongation primarily relies on intra-spindle pushing forces from kinesin-5 motors. The pathogenic yeast *Candida albicans* is unique because its spindle can elongate without a functional kinesin-5. Our study helps explain *C. albicans*’ diminished kinesin-5 requirement by showing that its dynein, Dyn1, collaborates with its kinesin-5, Kip1, to facilitate spindle elongation. Cells lacking Kip1 activity by either *KIP1* deletion or Kip1 inhibition have more Dyn1 and astral microtubules, indicating spindle elongation happens through enhanced pulling on spindle pole body-bound astral microtubules by cortical Dyn1. When Dyn1 activity is depleted, spindle elongation speed is normal, but anaphase spindles persist for extended periods, and the number and length of astral microtubules increases. Depletion of both motors prevents spindle elongation and is lethal. The significance of astral microtubules in cells lacking either motor is highlighted by the lethal effects of microtubule-destabilizing drugs and exposure to low temperatures (8°C) that compromise microtubule stability. These findings demonstrate that *C. albicans* spindle elongation involves cooperative forces from Kip1 and Dyn1 to ensure chromosome separation during cell division and that *C. albicans* adapts to changes in abundance of these motors through alterations in their astral microtubule content.

## Introduction

Mitotic chromosome segregation involves the equal partitioning of genetic information to daughter cells before cell division. This process depends on assembly of a bipolar microtubule (MT)-based structure called the mitotic spindle, which must form stable attachments to chromosomes, test that these connections were made correctly, and then separate sister chromatids to opposite ends of the dividing cell. All these activities require the help of teams of MT-binding motor proteins that apply directional forces on and within the spindle^1–3^. Spindles also rely on a fine-tuned balance of forces from MT polymerization and depolymerization events that are regulated by other MT-associated proteins^1–3^.

Physical separation of disjointed sister chromatids occurs during anaphase. In most eukaryotes, sister chromatids are first pulled towards the poles by depolymerizing the microtubules (MTs) bound to chromosomes, while overall spindle length is maintained. This process is referred to as anaphase A. In anaphase B, overlapping interpolar microtubules (ipMTs) in the spindle midzone slide apart, causing the two opposing spindle poles to separate and the disjoined chromatids to move toward their respective cellular compartments^4,5^. Surprisingly, a universal mechanism explaining the origins of all forces required for anaphase B is still lacking as research continues to uncover more organism-specific spindle elongation force-producers and regulatory processes thereof ^6^. In general, though, the major spindle forces of anaphase B typically include (i) interpolar microtubule (ipMT) sliding by spindle motors, (ii) ipMT braking by MT crosslinkers, (iii) pulling of astral microtubules (aMTs) from the cell cortex, and (iv) ipMT dynamics by MT plus-end polymerization and depolymerization, and (v) MT minus-end depolymerization manifesting as poleward flux^3,7–17^. Different combinations of these force-generating components generate distinct anaphase B mechanisms in different cell types.

In *Candida albicans* and other yeasts, MTs are attached directly to the SPBs and are stabilized, resulting in the absence of MT flux^18^. Therefore, anaphase A does not contribute to segregation of their chromosomes. Instead, chromosome segregation happens in anaphase B, where ipMT sliding by kinesin motor proteins in the spindle midzone elongate the spindle, contemporaneously pulling each sister chromatid poleward^19^. Here, MT midzone-localized plus-end directed kinesin-5 motors crosslink antiparallel ipMTs and slide them outward, past each other^9,15,17,19–23^. What remains unclear is the extent to which kinesin-5 pushing forces are required in spindle elongation as studies have shown that alternative or complementary mechanisms by motors located at the cell cortex may also contribute forces for anaphase spindle elongation^10,24–26^. For example, laser-mediated MT surgery experiments revealed that external forces are applied on the spindle via aMTs emanating from the spindle poles^10,25,27^. Here, dynein motors near the cell cortex bind these aMTs and apply pulling forces on the spindle^28–32^. In filamentous fungi, these dynein-mediated pulling events generate the primary spindle elongation force, while their kinesin-5 proteins are mainly employed to constrain the rate of spindle elongation^10,24^.

Previous studies in *C. albicans* showed that its dynein motor (Dyn1) is important in nuclear positioning, however, its role in spindle elongation was not defined^33^. Therefore, we studied the mitotic spindle phenotypes of cells depleted of Dyn1 as well as cells that lacked both Dyn1 and kinesin-5 (Kip1) activity. We observed that cells lacking Dyn1 could form anaphase spindles but often failed to orient the spindle across the mother-bud junction. Dyn1-depleted cells that succeeded in segregating their nuclei appeared to rely on Kip1 and polymerization of increased numbers of aMTs for compensatory forces. Conversely, cells lacking Kip1 contained an elevated abundance of dynein motors and aMTs. In the absence of both Dyn1 and Kip1 activity, cells arrested with a pre-anaphase spindle. Based on these findings, we propose that Kip1, Dyn1, and MT dynamics provide cooperative but spatially distinct spindle forces during anaphase.

## Results

### Dynein has overlapping functions with Kip1 in anaphase spindle elongation

In wild type *C. albicans*, the duration of anaphase is approximately 20 minutes. When Kip1 function is lost by *KIP1* gene deletion, bipolar spindles can form and elongate but are ∼25% shorter^34^. *kip1*Δ/Δ spindles also have drastically longer and more numerous aMTs attached to their poles. A possible explanation for these observations is that *kip1*Δ/Δ cells rely on Dyn1 motors distributed near the cell cortex to bind these aMTs and pull on them to elongate the spindle, thereby compensating for the loss of intra-spindle pushing forces from Kip1. If this is the case, Kip1 and Dyn1 likely function together during anaphase in wild type cells. To investigate this putative functional overlap between Kip1 and Dyn1 in spindle elongation, we studied the growth and mitotic spindle phenotype of cells in which activity one or both motors was eliminated.

On solid YPD media, growth of *dyn1*Δ/Δ cells at 30 °C was reduced compared to wild type but was comparable to cells lacking either *KIP1* or the kinesin-14 *KAR3* (**Figure 1A**). On YPD media supplemented with 100 µM of the Kip1-specific inhibitor aminobenzothiazole (ABT)^35^, *dyn1*Δ/Δ cells were not viable. Similar growth defects were observed when *dyn1*Δ/Δ cells were grown in liquid cultures supplemented with 100 µM ABT (**Figure 1B**). To identify cell cycle defects that could explain the death of ABT-treated *dyn1*Δ/Δ cells, we generated wild type and *dyn1*Δ/Δ cells that expressed β-tubulin fused to green fluorescent protein (Tub2-GFP) and mScarlet-labeled nucleolar protein (Nop1-mScarlet), allowing us to track morphologic changes in their mitotic spindle and nucleus, respectively, during progression through mitosis. We then cultured these cells in ABT-free media and then added 50 µM ABT for 3 h before imaging them by time-lapse fluorescence microscopy. **Figure 1C** shows that, following incubation with ABT, *dyn1*Δ/Δ cells contained a single spot of nuclear fluorescence and a short, immobile pre-anaphase spindle that persisted in the mother cell compartment for the entire time-lapse even after the bud cell had grown to its typical size (over 200 minutes). In comparison, untreated wild type cells completed anaphase within 20 minutes. Suspecting that ABT addition prevented anaphase initiation, we added increasing concentrations of ABT to unsynchronized cell populations of *dyn1*Δ/Δ mutants and compared the lengths of their spindles to those of wild type and *dyn1*Δ/Δ cells that were not exposed to ABT. We observed that, as ABT concentration increased, the average spindle lengths in *dyn1*Δ/Δ cells decreased (mean lengths: *wild type* = 2.26 µm, *dyn1*Δ/Δ = 2.54 µm, *dyn1*Δ/Δ + 75 µM ABT = 1.4 µm, *dyn1*Δ/Δ + 100 µM ABT = 1.0 µm; P<0.05) and that very few ABT-treated *dyn1*Δ/Δ cells had spindles longer than 3 µm (**Figure 1D**). These data show that cells lacking both Kip1 and Dyn1 activity cannot enter anaphase, indicating that both motors contribute forces for spindle elongation.

**Figure 1.**
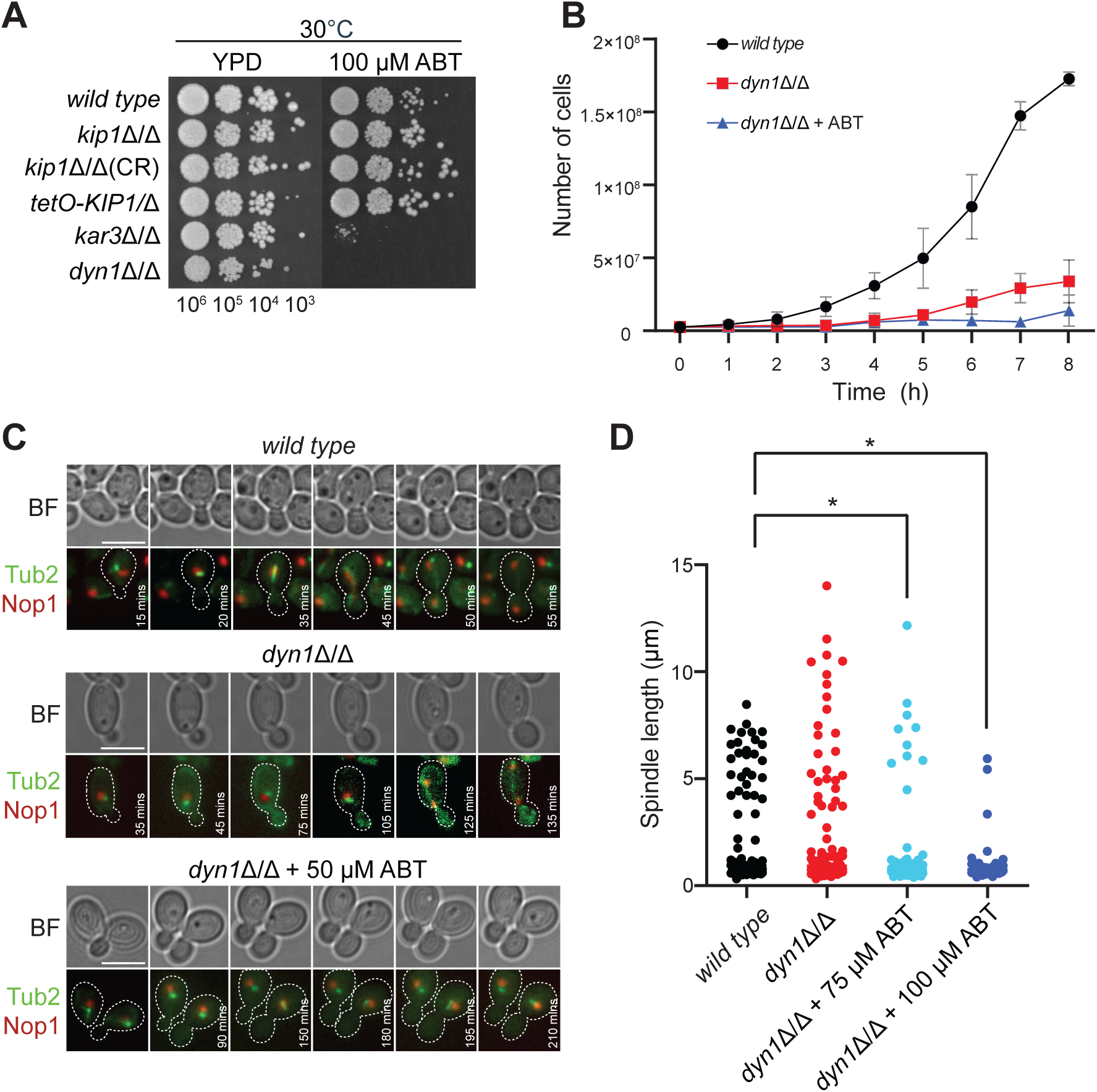
Simultaneous loss of Dyn1 and Kip1 function is lethal. (A) Wild type (CF027), *kip1*Δ/Δ (CF311, CF429, CF436), *kar3*Δ/Δ (CF024) *dyn1*Δ/Δ (CF495) cells were plated on YPD or YPD + 100 µM ABT. Cells were serially diluted to the indicated concentrations and 5 µL drops were plated and incubated for 2 days at 30°C. (B) Cell growth assay of wild type (CF027), *kip1*Δ/Δ (CF311) and *dyn1*Δ/Δ (CF495) with and without 100 µM ABT. Strains in YPD media were diluted 2.5 x 10^6^ cells per mL, incubated at 30°C and counted every hour using a hemocytometer. Data points represent an average from three independent experiments +/- standard deviation (SD). (C) Time-lapse microscopy of wild type and *dyn1*Δ/Δ cells expressing Nop1-mScarlet and Tub2-GFP (CF417, CF486). ABT (50 µM) was added to *dyn1*Δ/Δ cells and incubated for 3 h before imaging. Images were collected every 5 mins with exposure times of 150 ms for both fluorescent tags. Scale bar, 5 µm. (D) Quantification of all spindle lengths observed in unsynchronized cell populations of wild type (CF405) (n=101), *dyn1*Δ/Δ (CF542) (n=101), *dyn1*Δ/Δ + 75 µM ABT (n=101), and *dyn1*Δ/Δ + 100 µM ABT (n=50) cells (*P <* 0.05, Student’s t-test). Cells were grown to mid-logarithmic phase in SDC-sucrose + 20 % galactose and ABT was then added and incubated for 1 h.

### Dynein localization on astral MTs increases in the absence of Kip1

To further investigate the involvement of Dyn1 in anaphase spindle elongation in *C. albicans*, we constructed a Dyn1-mNeon-labeled strain by inserting the coding sequence of the fluorescent protein mNeon in frame with the 3′ end of the chromosomal gene for dynein heavy chain *DYN1* (Dyn1-mNeon). We engineered this strain to co-express a Tub2-mScarlet fusion protein, allowing us to compare Dyn1 localization in relation to the mitotic spindle and aMTs. In 70% of cells with small buds and metaphase spindles, Dyn1-mNeon fluorescence was prominent on the plus ends of aMTs that extended towards or into the bud (**Figure 2A, *wild type* panel, row 1**). This asymmetric localization of Dyn1 has been observed in other *C. albicans* studies, and in *S. cerevisiae*, and helps with spindle positioning and orientation, ensuring that the daughter SPB enters the bud during cell division^31,36,37^. In 82% of cells with long anaphase spindles, in which the daughter SPB had moved into the bud, two to four Dyn1-mNeon spots were visible at the plus ends of aMTs from both SPBs (**Figure 2A, *wild type* panel, rows 2 and 3**). In these cells, Dyn1-mNeon was usually localized near the cortex of the mother and daughter compartments.

**Figure 2.**
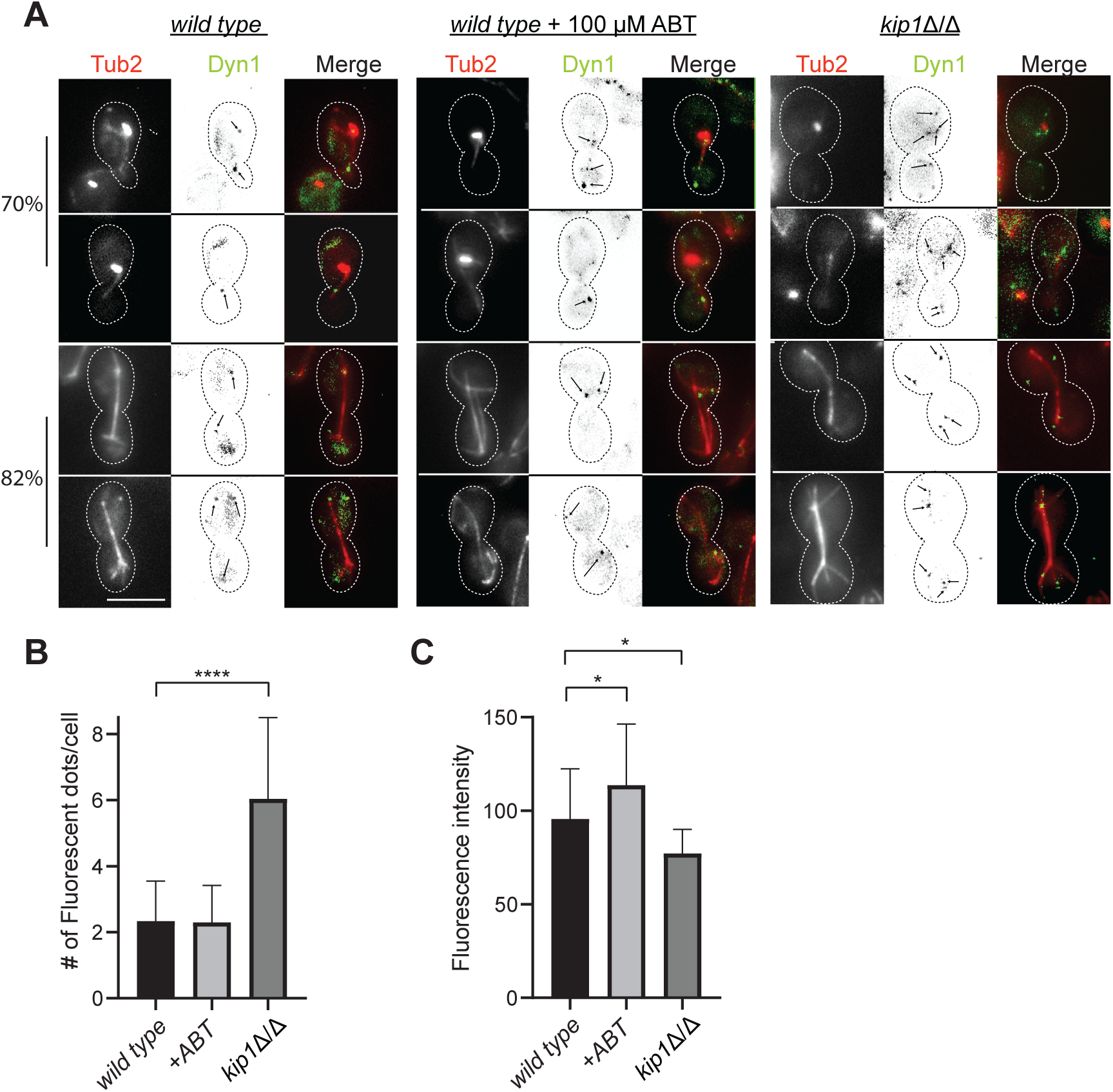
Dyn1 localization and abundance increases when Kip1 function is lost or inhibited. (A) Images of wild type and *kip1*Δ/Δ cells expressing Tub2-mScarlet and Dyn1-mNeon (strains CF546, CF543). For Kip1 inhibition in wild type cells, 100 µM of ABT was added and incubated for 3 h. Representative cells at different stages of mitosis were selected. All cells were obtained from logarithmically growing, unsynchronized cultures in SDC-sucrose medium at 30°C. Scale bars, 5 µm. (B) The number of Dyn1 fluorescent patches in cell types from (A) was counted and graphed (n = 30 for each strain). (C) Quantification of Dyn1 fluorescence intensity in all three cell types from (A). Dyn1 fluorescence intensity was measured against background using ImageJ Software.

If Dyn1 provides pulling forces on the aMTs of each SPB for anaphase spindle elongation, we speculated that the abundance of Dyn1 motors on these MTs would increase when Kip1 pushing forces from within the spindle were eliminated or inhibited. To investigate this possibility, we examined the location and abundance of Dyn1-mNeon in *kip1*Δ/Δ and ABT-treated wild type cells. Compared to wild type cells, *kip1*Δ/Δ mutants usually exhibited two or three more fluorescent Dyn1 foci in both the mother and daughter cell, regardless of the cell cycle phase (**Figure 2A and 2B)** (average number of Dyn1-mNeon foci: *wild type* = 2, *kip1*Δ/Δ = 6, ABT-treated *wild type* = 2). Moreover, there was a 1.2-fold increase in the overall Dyn1-mNeon fluorescence intensity in ABT-treated wild type cells compared to untreated cells (average Dyn1 intensity: *wild type* = 95.6, *kip1*Δ/Δ = 77.2, ABT-treated *wild type* = 113.7) (**Figure 2A and 2C**). We propose that this increase in Dyn1 fluorescence with ABT treatment is an indication that even more Dyn1 is needed to compensate for ABT’s inhibitory effect on Kip1. Rather than eliminating pushing forces by Kip1 motors within the spindle, ABT-inhibited Kip1 forms a rigor-like complex with the MT^35^. Thus, a considerable increase in pulling forces from cortical Dyn1 may be required to break apart a rigid bipolar spindle stabilized by ipMT-locked Kip1 motors.

### Anaphase duration is nearly tripled in *dyn1*Δ/Δ cells

To understand how Dyn1 contributes to spindle elongation in *C. albicans*, we tracked spindle dynamics and nuclear segregation in dividing *dyn1*Δ/Δ cells using time-lapse imaging of Tub2-GFP and Nop1-mScarlet fluorescence, respectively. We found that most *dyn1*Δ/Δ cells formed normal-looking bipolar spindles that elongated and separated the Nop1-mScarlet fluorescence into two distinct foci. However, only 30% (32.3%) of these cells completed anaphase at the same rate as wild type (avg anaphase time; *wild type* = 21.3 mins, *dyn1*Δ/Δ, bottom, pink cluster = 25.9 mins) (**Figure 3A**). The rest of these *dyn1*Δ/Δ cells took considerably longer (*dyn1*Δ/Δ, top, blue cluster = 34.96 +/- 4.64 mins). When we made kymographs of the wild type and *dyn1*Δ/Δ spindles, beginning from bipolar spindle appearance and ending at late anaphase, we found that 60% of *dyn1*Δ/Δ cells remained in anaphase for an average of 60 mins before the spindle disassembled (P<0.0001) (**Figure 3B**). Just over half of these cells (∼60%) underwent anaphase spindle elongation within the mother cell (**Figure 3C and 3E**). The rest (39.4%) succeeded in elongating the spindle across the mother-bud junction (**Figure 3D, cell 2, and Figure 3E and 3F**). Thus, even the *dyn1*Δ/Δ cells that succeeded in moving the daughter SPB into the bud were delayed in anaphase completion and spindle disassembly. Another curious finding was that many *dyn1*Δ/Δ anaphase spindles showed signs of buckling, especially when these spindles were trapped in the mother cell compartment (**Figure 3D, cell 1**). These data demonstrate that *dyn1*Δ/Δ cells have difficulty positioning the spindle for entry into the mother-bud junction and that Kip1 provides the bulk, but not all, of the force for spindle elongation. We suspect that the spindle buckling events in *dyn1*Δ/Δ cells result from an inability of intra-spindle pushing forces from Kip1 to overcome the compression forces on the spindle when Dyn1 is absent^38^.

**Figure 3.**
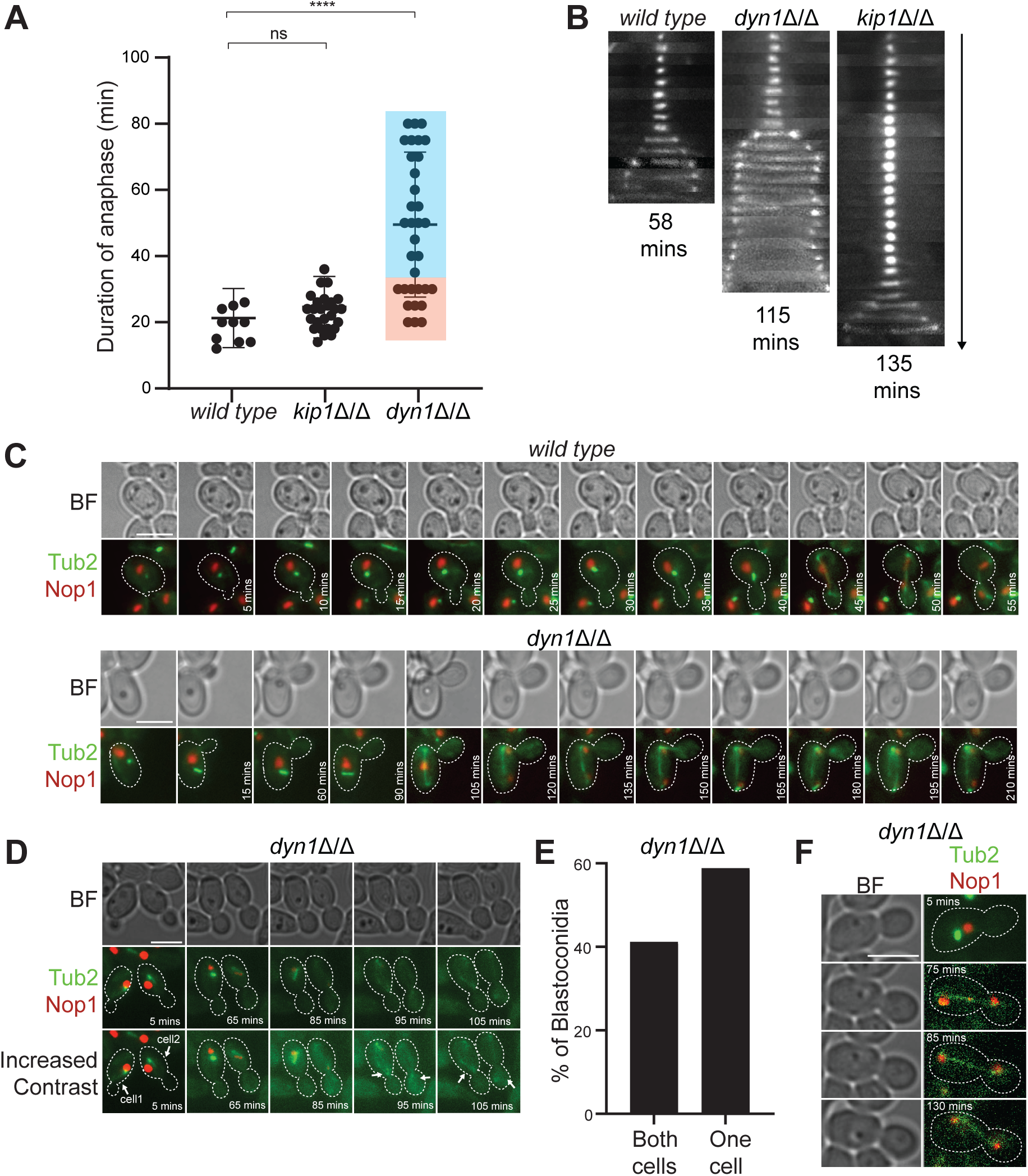
Dyn1 mutants exhibit unusual anaphase dynamics. (A and B) Quantification of wild type, *kip1*Δ/Δ, and *dyn1*Δ/Δ cells using time-lapse microscopy to analyze the duration of anaphase. Long (2 to 4 h) time-lapse series were captured with 150-ms exposures to measure the length of time from start to the end of anaphase for Tub2-GFP (n = 11), Tub2-GFP *kip1*Δ/Δ (n = 28), and Tub2-GFP *dyn1*Δ/Δ (n = 34) (*P* < 0001, Student’s t-test). Representative kymograph is illustrated in (B) for all three strains. Kymograph for *kip1*Δ/Δ was previously published^34^. (C and D) Time-lapse microscopy of wild type and *dyn1*Δ/Δ cells expressing Nop1-mScarlet and Tub2-GFP (CF417, CF486). Images in (C) illustrate anaphase events occurring in mother cell only. Images in (D) represent two scenarios of anaphase events in *dyn1*Δ/Δ cells. In Cell 1, anaphase is occurring within the mother compartment and shows ‘buckling’ of the anaphase spindle Cell 2 shows normal anaphase. Images were collected every 5 mins with exposure times of 150 ms for both fluorescent tags. Scale bar, 5 µm. (E) Quantification of anaphase events in *dyn1*Δ/Δ cells occurring across mother-bud junction or solely in the mother cell. (F) Time-lapse microscopy of *dyn1*Δ/Δ cells expressing Nop1-mScarlet and Tub2-GFP (CF486). Spindle disassembly is not initiated even after the spindle has elongated into the bud.

### MT polymerization forces facilitate *C. albicans* spindle positioning and elongation

Previous studies in *S. pombe* showed that MT polymerization-derived pushing forces are sufficient to promote spindle pole separation and assembly of a bipolar spindle in the absence of molecular motors^2,39,40^. Similar results have been observed in higher eukaryotes, signifying a functional conservation of MT-based forces within the spindle^41,42^. A curious phenotype of our *dyn1*Δ/Δ cells was that they formed longer and more numerous aMTs than wild type cells (mean aMT lengths: *wild type* = 2.2 µm ± 1.25 SD, *dyn1*Δ/Δ = 3.9 µm ± 2.8 SD; mean aMT number: *wild type* = 1 aMT ± 1 SD, *dyn1*Δ/Δ = 4 ± 2 SD), suggesting that *C. albicans* may use MT polymerization forces in addition to motor-derived forces for spindle morphogenesis and chromosome segregation (**Figure 4A, 4B, and 4C**). In addition, aMT dynamics could provide pushing forces to orient the spindle in the absence of dynein^3^. To investigate this possibility, we studied *dyn1*Δ/Δ cell growth under two distinct MT polymerization-disrupting conditions: (1) Nocodazole treatment and (2) cold temperature (8°C). As expected, when we incubated liquid cultures of unsynchronized wild type and *dyn1*Δ/Δ cells with 50 µM nocodazole for 6 h and then obtained snapshots of the spindles in these cell populations by fluorescence microscopy, we found that both strains formed normal-looking mitotic spindles but lacked aMTs (**Figure 4D**). In a dilution spot assay of cells grown on solid YPD media supplemented with 100 µM Nocodazole, *dyn1*Δ/Δ cells showed the strongest growth defect (**Figure 4E**). Likewise, *dyn1*Δ/Δ cells were much more sensitive to Nocodazole compared to wild type and *kip1*Δ/Δ cells when grown in liquid cultures (**Figure 4F**).

**Figure 4.**
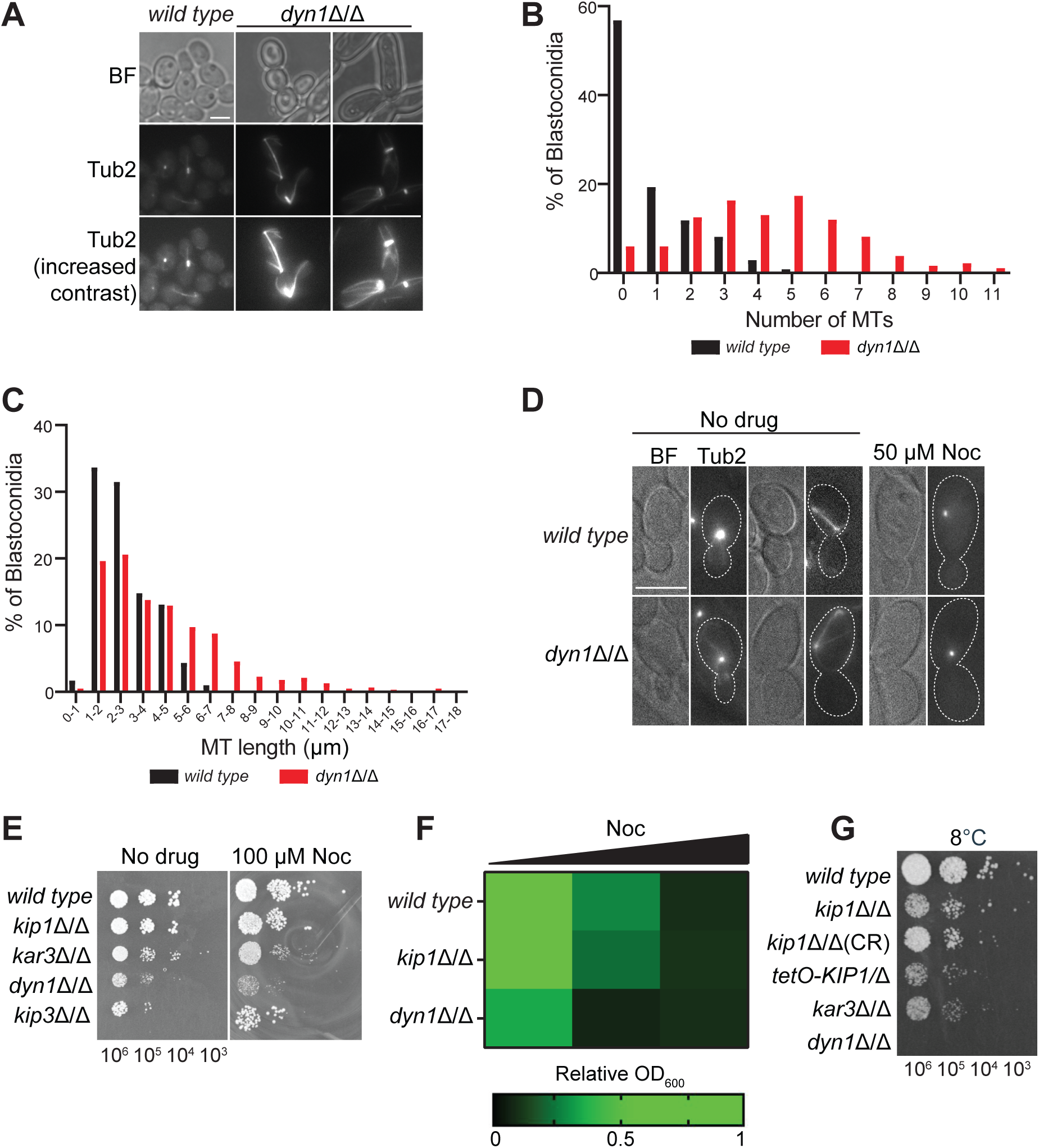
*dyn1*Δ/Δ cells have longer and more numerous aMTs. (A) Representative images of wild type and *dyn1*Δ/Δ cells expressing Tub2-GFP to illustrate the difference in aMT number and lengths. (B) The number of aMTs in wild type (CF405) and *dyn1*Δ/Δ (CF488) expressing Tub2-GFP was counted in cells with visible spindles. (C) For cells in panel B that contained aMTs, aMT length was determined by measuring the distance between the metaphase spindle pole and the plus end. These lengths were organized into bins of the size ranges indicated. (wild type: n = 413; *dyn1*Δ/Δ: n = 617) (D) Static images of wild type (CF405) and *dyn1*Δ/Δ cells (CF488) expressing Tub2-GFP in SDC-sucrose and galactose. 50 µM of Nocodazole was added and cells were incubated for 6 h and imaged. Scale bar, 5 µm. (E) Wild type (CF027), *kip1*Δ/Δ (CF311), *kar3*Δ/Δ (CF024) *dyn1*Δ/Δ (CF495), *kip3*Δ/Δ (CF254) cells were plated on YPD or YPD + 100 µM Nocodazole. Cells were serially diluted to the indicated concentrations and 3 µL drops were plated and incubated for 2 days at 30°C. (F) Cell growth assay of wild type (CF027), *kip1*Δ/Δ (CF311) and *dyn1*Δ/Δ (CF495) with increasing concentrations of Nocodazole (100-250 µM). Strains in YPD media were diluted 0.2 optical density (OD) cells per mL, incubated at 30°C, and OD was measured every hour. (G) Wild type (CF027), *kip1*Δ/Δ (CF311, CF429, CF436), *kar3*Δ/Δ (CF024) *dyn1*Δ/Δ (CF495) cells were plated on YPD. Cells were serially diluted to the indicated concentrations and 5 µL drops were plated and incubated for 25 days at 8°C.

Similar growth defects were observed when wild type, *kip1*Δ/Δ, *kar3*Δ/Δ and *dyn1*Δ/Δ cells were spotted on YPD plates and grown at a cold temperature (8°C) (**Figure 4G**). At this temperature, MT polymerization kinetics are dramatically slower and MTs are less stable^43^. These results suggest that *C. albicans* uses MT polymerization forces to help in spindle positioning, and that these cells can employ these MT forces when faced with a deficit or imbalance of motor protein-generated pushing or pulling forces.

## Discussion

It is well-established that pulling forces on the plus ends of SPB-bound aMTs by cortical Dyn1 motors are important for proper mitotic spindle orientation and nuclear migration in *C. albicans*^33,36^. Our observations support this but show that Dyn1 is not essential for spindle positioning. In the absence of Dyn1, aMTs are longer and more abundant. Loss of aMTs by nocodazole addition or cold temperatures was lethal to *dyn1*Δ/Δ cells, but not to wild type or even *kip1*Δ/Δ cells. These observations indicate collaborative action between dynein and aMT dynamics in spindle positioning. We propose that the correct spindle positioning we observed in cultures of Dyn1-deficient *C. albicans* cells is most likely enabled by chance interactions between dynamic aMTs interacting with the cell cortex that guide movement of the daughter SPB toward and through the mother-bud junction^33^.

The experiments described here also demonstrate that Dyn1 has the added function of working collaboratively with Kip1 to facilitate spindle elongation in *C. albicans*. One piece of supporting evidence is that simultaneous loss of Kip1 and Dyn1 activity was lethal and appears to stem from the cell’s inability to initiate anaphase. Another is that when Kip1 activity was chemically inhibited or eliminated by *KIP1* deletion, Dyn1 motors and aMTs became more abundant. We presume that the additional numbers of Dyn1 and aMTs afford increased pulling force from the cell cortex to compensate for loss of outward pushing forces from the spindle midzone, helping anaphase spindle elongation to occur, albeit much more slowly than in the presence of Kip1. When Dyn1 activity was eliminated by *DYN1* deletion, we observed that spindles usually elongated at rates similar to wild type *C. albicans* cells, indicating that Kip1 provides most of the force needed for spindle elongation via ipMT sliding at the spindle midzone. However, it is also possible that *C. albicans* modifies mitotic spindle force-balance mechanisms in lieu of missing motors by changing the abundance or activity of other motors that contribute to antiparallel sliding of ipMTs or that alter the dynamics of MT plus ends in the midzone. Notably, the kinesin-8 motor (Kip3) in *C. albicans* has been shown to influence spindle length throughout mitosis via its microtubule-depolymerizing activity^44^. In *S. cerevisiae*, a complex interplay exists between MT depolymerization by kinesin-8 and ipMT sliding activity that contributes to spindle elongation and disassembly^45,46^.

Similar to previous observations by Findley *et al*. 2008^36^, we found that *dyn1*Δ/Δ cells remained in anaphase for up to three times longer than wild type. Although this delay in anaphase completion was often due to mislocalization of the spindle into the mother cell, which is reflective of Dyn1’s role in spindle positioning, many of the *dyn1*Δ/Δ cells that succeeded in elongating the spindle across the mother-bud junction also contained persistent anaphase spindles. These findings suggest that dynein activity is important for assisting in spindle elongation after anaphase onset and that the protracted anaphase of *dyn1*Δ/Δ yeast cells cannot be fully accounted for by activation of a Bub2p-mediated checkpoint that inhibits spindle disassembly until the daughter SPB and daughter nucleus enter the daughter cell^36,47^.

Although research continues to demonstrate that conserved motor proteins can have varying levels of involvement in essential cellular functions for different eukaryotes, harmonization of kinesin-5 and dynein activities are clearly a consistent feature of mitotic spindle dynamics across eukaryotes. In organisms where kinesin-5 activity is essential for viability, spindle elongation is predominantly mediated by ipMT sliding by these motors at the spindle midzone^22^. For instance, *Drosophila* embryos use ipMT sliding by their kinesin 5 (Klp61F) coupled to suppression of MT disassembly at the poles to push spindle poles apart^17,48^. *S. cerevisiae* cells employ two kinesin-5s (Cin8 and Kip1) to slide ipMTs apart^19,49^. Although it has been reported that Dyn1 also facilitates spindle elongation in *S. cerevisiae*, the extent of its pulling force during this event is unclear compared to the pushing forces generated by both kinesin-5 motors^21^. In contrast, kinesin-5 is not essential in *Caenorhabditis elegans* and *Dictyostelium discoideum*^8,50^. In these eukaryotes, external pulling forces from aMT depolymerization and dynein motility are dominant, while the role of their respective kinesin-5 motors is to restrict ipMT sliding at the spindle midzone. In the filamentous fungi, *U. maydis*, although kinesin-5 is not essential, it is important in the slow phase of anaphase, while dynein contributes to the fast phase^24^. Similarly, we show that *C. albicans* employs the activities of both dynein and kinesin-5 for spindle elongation.

Why *C. albicans* and *U. maydis* rely more heavily on dynein for spindle elongation than *S. cerevisiae* and some other fungi is an interesting question. An answer may relate to differences in the sizes and shapes of their cells, and the distances over which the nucleus must be moved. *C. albicans* is a polymorphic fungus that can switch between yeast, pseudohyphae, and hyphal growth forms^51^. Dyn1’s function in spindle elongation is likely more pronounced in hyphal spindles as the spindle lengths in these cells exceed that of spindles of yeast cells^52^. Here, involvement of pulling forces on MTs would be the preferred mechanism of moving objects greater than 10 µm, as MT pushing could experience a high rate of MT buckling^3^. Thus, it is plausible that dynein and kinesin-5 motors are actively regulated to provide different degrees of spindle elongation forces according to the size and growth form of *C. albicans.* Further investigations of *C. albicans* motors are required to test these theories. Studies like this could also show how this fungus may be able regulate its mitotic motors to asymmetrically distribute its genetic information and generate aneuploidies that foster its adaptability to human hosts and its resistance to antifungal drugs^53^

## Materials and Methods

### *C. albicans* Strains and Genetic Manipulations

A list of *C. albicans* strains used in this study is presented as **Table 1**. For a list of oligonucleotides used in *C. albicans* strain construction, please refer to **Table 2**. Gene disruption of the *C. albicans DYN1* open reading frame (Candida Genome Database: *orf19.5999*) was conducted by transformation and integration of a linear cassette containing a selectable marker. PCR amplification was used to generate disruption cassettes where a selectable marker was flanked by approximately 50 bp of *C. albicans* genomic sequence immediately 5’ and 3’ of the *DYN1^+^* coding region. Disruption of *DYN1^+^* in a wild type background (CF027) was conducted sequentially. First a *dyn1::LEU2^+^* cassette was amplified from *pSN40* ^54^ using P304 and P305 and transformed into CF027. Correct *dyn1::LEU2^+^* cassette integration was confirmed using primers P306 and P13. Second, a *dyn1::HIS1^+^* cassette was amplified from *pSN52* ^54^ using primers P304 and P305 and transformed into *dyn1::LEU2* to create CF495. Integration of the disruption cassette at the correct location was confirmed by PCR amplification across the junctions of integration using primers P306 and P12.

**Table 1.**
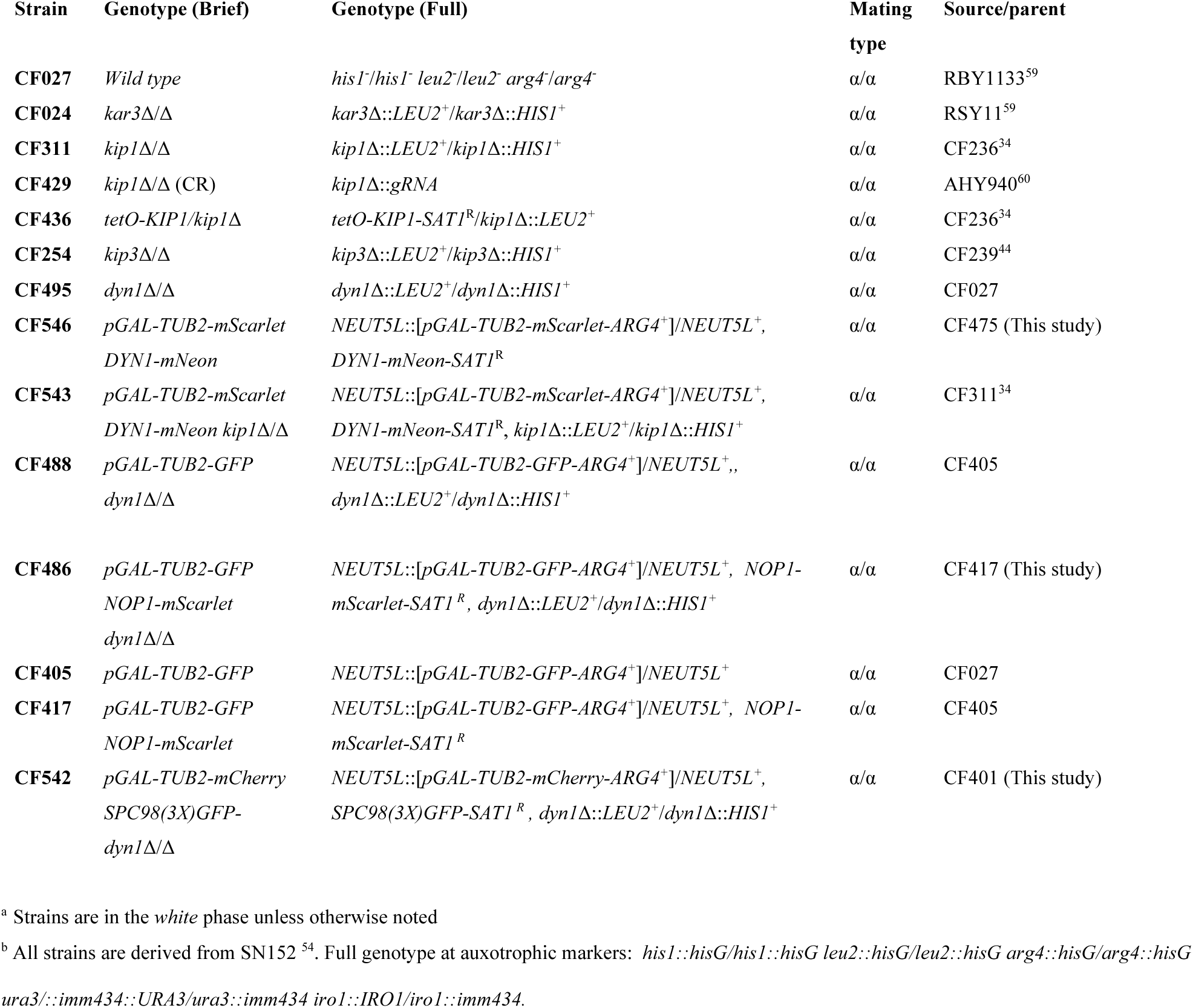
Names, genotypes, mating types, and sources of the strains used in this study.

**Table 2.**
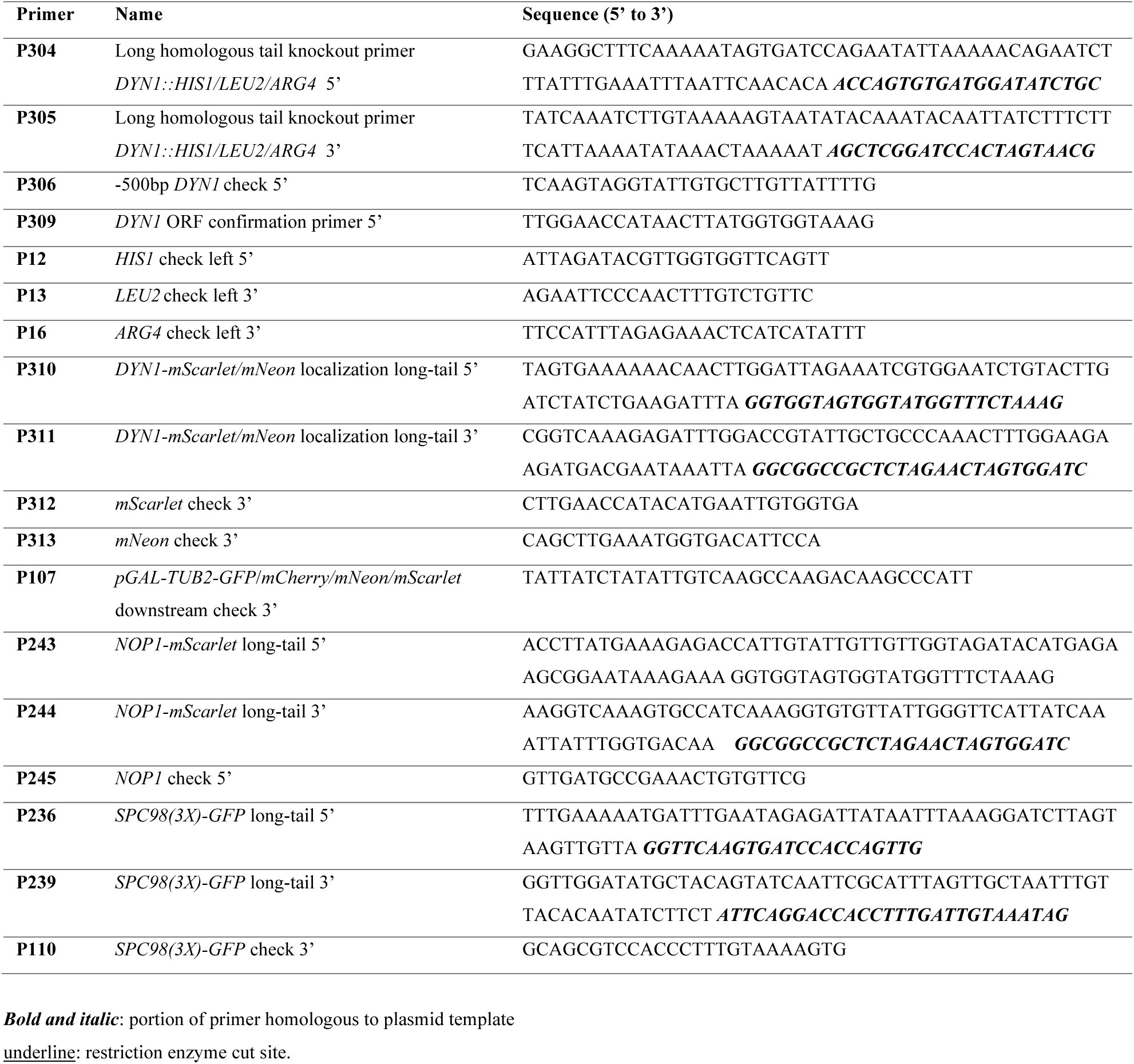
Oligonucleotide primers used in strain construction.

Strains expressing fluorescently labeled β-tubulin were constructed using the plasmid pGAL:TUB2-GFP-NEUT5L and pGAL:TUB2-mScarlet-NEUT5L and linearized by *KPN1* to target to the neutral NEUT5L locus, as previously described^55^. Correct integration for pGAL vectors was confirmed by PCR using primers P16 and P107. To visualize the nucleus, the nucleolar protein Nop1 was fluorescently labeled using the plasmid *pmScarlet-SAT1* (kindly provided by Dr. Richard Bennet) and amplified by primers P243 and P244 ^56^. Correct integration was confirmed using P312 and P245. Strains expressing fluorescently labeled spindle pole body component, SPC98, were constructed using a pGFP(3X) with primers P236 and P239. Correct integration as confirmed using primers P107 and P110.

Fluorescent tagging of *DYN1^+^* in a wild type and *kip1*Δ/Δ background was accomplished using the method described by Gerami-Nejad *et al*.^57^. Briefly, long-tailed primers P310 and P311 and the plasmids *pmNeon-SAT1* or *pmScarlet-SAT1* as a template to create an integration cassette bearing approximately 50 base pairs of *DYN1^+^* ORF immediately before the stop codon, and of sequence 3’ to the ORF. Correct integration was confirmed by PCR using primers P309, P312 and P313.

### *C. albicans* Transformation

*C. albicans* transformations were done using the lithium acetate-polyethylene glycol (PEG) heat shock method as previously described with minor modifications^58^. Transformations involving selection using the SAT1 gene were recovered in YPD (1% yeast extract, 2% peptone and 2% glucose) at 30°C for 4 h to allow expression of the ClonNAT resistance gene before plating on Nourseothricin selection medium.

### *C. albicans* Growth Media and Chemical Reagents

All strains were maintained on YPD plates. YPD was supplemented with 200 µg/ml nourseothricin (clonNat; Werner BioAgents) for selection of positive SAT1 gene integration. Selection for auxotrophic markers was conducted using synthetic dropout (SD) medium containing 0.66% yeast nitrogen base, 0.2% yeast dropout mix lacking uracil, arginine, leucine, and histidine, 2% glucose, and 200 mg/L uridine and supplemented with 200 mg/L histidine, leucine, and/or arginine where required. Experimental cultures were grown to mid-logarithmic phase in YPD unless otherwise indicated. To create dilutions for spot assays, logarithmically growing cells were diluted to 1.0 x 10^6^ cells/ml in phosphate-buffered saline (PBS). Serial dilutions of 10^5^,10^4^, and 10^3^ cells/ml were made. Five µL of cell culture dilutions was pipetted for each spot, and plates were incubated at 30°C for 2 days, unless otherwise indicated.

### Microscopic Imaging of Spindle and Nuclear Morphologies and Motor Localization

Static images of cells were captured using a Zeiss Axio Observer epifluorescence microscope with a 100X (1.40 NA) oil objective AxioCam hRM camera controlled by Axiovision software. Time-lapse imaging and some static images were recorded using the Olympus IX83 with a 100X oil objective (1.4 NA), and Andor Zyla 4.2 Plus camera controlled by the cellSens software. For time-lapse and static imaging, logarithmically growing cells were immobilized between an agarose pad and a glass coverslip, as previously described^55^. For time-lapses, images were captured in five z-slices 0.8 µm apart. Spindle length measurements were analyzed using ImageJ (NIH). Analysis was done with GraphPad Software. Figures were constructed in Adobe Photoshop and Adobe Illustrator.

## Acknowledgements

We are grateful to the efforts at the *Candida* Genome Database (http://www.candidagenome.org/) for archiving and annotating *Candida* sequence information. We thank Peter Davies (Queen’s University) for the use of their microscopy facilities. We thank Richard Bennett and Corey Frazer (Brown University) for the mNeon- and mScarlet-encoding genes. We thank Leah Cowen (University of Toronto) for providing the plasmid used to generate the tetracycline-regulatable (TR) promoter system. This work was supported by a National Sciences and Engineering Council of Canada grant (RGPIN-2019-05924) and a Canadian Institutes of Health Research grant (PJT- 169149). J.S.A. is a former Canada Research Chair (Tier 2) in Structural Biology and an Ontario Early Researcher Award recipient. I.S. is an Ontario Graduate Scholarship recipient.

